# Fast-scBatch: Batch Effect Correction Using Neural Network-Driven Distance Matrix Adjustment

**DOI:** 10.1101/2024.06.25.600557

**Authors:** Fu Chen, Leqi Tian, Teng Fei, Tianwei Yu

**Affiliations:** School of Data Science, The Chinese University of Hong, Kong, Shenzhen (CUHK-Shenzhen), Shenzhen, Guangdong, China; School of Data Science, The Chinese University of Hong, Kong, Shenzhen (CUHK-Shenzhen), Shenzhen Research, Institute of Big Data, Shenzhen, Guangdong, China; Memorial Sloan Kettering Cancer Center, New York, New York, USA

**Keywords:** correlation structure, batch effect, principal component analysis, deep learning, single-cell sequencing

## Abstract

Batch effect is a frequent challenge in deep sequencing data analysis that can lead to misleading conclusions. Existing methods do not correct batch effects satisfactorily, especially with single-cell RNA sequencing (scRNA-seq) data. To address this challenge, we introduce fast-scBatch, a novel and efficient two-phase algorithm for batch-effect correction in scRNA-seq data, designed to handle non-linear and complex batch effects. Specifically, this method utilizes the inherent correlation structure of the data for batch effect correction and employs a neural network to expedite the process. It outputs a corrected expression matrix, facilitating downstream analyses. We validated fast-scBatch through simulation studies and on two scRNA-seq datasets, demonstrating its superior performance in batch-effect correction compared to current methods, as evidenced by visualization using UMAP plots, and metrics including Adjusted Rand Index (ARI) and Adjusted Mutual Information (AMI).

## 1 INTRODUCTION

In recent years, technological advancements have significantly enhanced our ability to generate high-throughput single-cell gene expression data. However, due to limitations in sequencing technology, sample collection, and operational considerations, single-cell data are often generated in multiple experiments. These experiments can introduce batch-specific systematic variations, known as “batch effects” [1]. These differences can alter the statistical distribution of the gene count matrix and obscure the biological specificity we are interested in, leading to misinterpretation of cell types and cell heterogeneity, and consequently misleading scientific discoveries in downstream analyses [2]. Therefore, addressing batch effects is crucial for ensuring the fidelity of biological insights derived from the data. Appropriate batch effect correction methods can integrate experimental results accurately and interpretably, thereby enhancing the reliability and robustness of downstream analyses.

Various methods have been developed in recent years to correct batch effects, each following different strategies. For example, ComBat uses Bayesian methods to normalize data and reduce between-batch shifts and scaling [3]. Limma introduces linear models to detect and eliminate batch effects [4]. MultiCCA uses canonical correlation analysis (CCA) to capture data features and project them into subspaces to find correlations between different batches [5]. These methods are generally classified as linear modeling approaches and handle linear or log-linear batch effects well. They assume that batch effects are additive or multiplicative biases that can be corrected through linear transformations. However, as singlecell RNA sequencing (scRNA-seq) datasets become more complex and heterogeneous, batch effects become more intricate and non-linear, making linear model approaches less suitable.

To address these limitations, researchers have developed various nonlinear batch effect correction methods. For instance, the Mutual Nearest Neighbors (MNN) algorithm uses cell neighborhood relationships to eliminate batch effects [6]. This method identifies cell mappings between batches and establishes links by identifying mutual nearest neighbors, using these as “anchors” for alignment. Harmony is an iterative method that aligns datasets in latent space to remove batch effects, ensuring consistent distribution across integrated datasets [7]. Scanorama is a batch correction method based on neighborhood search, identifying common neighborhoods across batches in high-dimensional space for correction [8]. With the rapid advancement of deep learning technologies, many deep learning-based methods have also been applied to batch effect correction. For example, scVI (single-cell Variational Inference) uses variational autoencoders (VAE) to model single-cell gene expression data and correct batch effects in latent space [9]. HDMC (Hierarchical Distribution Matching and Contrastive Learning) employs high-dimensional manifold learning techniques to discover lowdimensional data representations, correcting batch effects in this representation [10]. The scDML method uses deep networks to embed samples into low-dimensional space, eliminating batch effects during the embedding process [11]. These methods leverage the powerful representation capabilities of deep learning to better capture and correct complex nonlinear batch effects.

We previously proposed scBatch, a two-phase batch-effect correction method [12]. It first uses QuantNorm [13] to obtain a corrected correlation matrix by quantile normalization, then uses a coordinate descent algorithm to restore the corrected count matrix to the original space. Despite its accuracy and convenience, the process is time-consuming on large datasets. With the increasing demand for processing large datasets, there is an urgent need to improve efficiency.

Here we propose a new method called fast-scBatch to correct batch effects. It bears some resemblance to scBatch in that it also uses a two-phase approach, and starts with the corrected correlation matrix in phase 1. On the other hand, the second phase of restoring the count matrix is newly designed to incorporate the idea of using dominant latent space [14] in batch effect removal, and a customized gradient descent-supported algorithm. Additionally, we explored supplementary algorithms that, while possibly less accurate, can compute faster in specific scenarios. We evaluated our method using simulation studies with data generated using the R package splatter [15], as well as two real scRNA-seq datasets, using UMAP plots and metrics such as Adjusted Rand Index (ARI) and Adjusted Mutual Information (AMI) to compare its performance with stateof-the-art methods.

## 2 METHODS

Fast-scBatch basically assumes a scenario where all cell types are present in each batch, and their proportions do not dramatically differ across batches. This assumption aligns with widely adopted practices in batch effect correction evaluations for their practicality and impartiality. Within this framework, we assess the correlation between cell count matrices within each batch and derive an expected correlation matrix devoid of batch effects. Subsequently, we utilize this corrected correlation matrix to obtain the batch-effect-free cell count data as desired.

### 2.1 Problem setup

Fast-scBatch requires the information of th *p*×*n* cell count matrix, where *n* is the total number of cell samples in all batches and *p* is the number of variables (e.g. genes). The batch information, i.e. the batch annotation for each sample, is also needed. The size of samples in each batch is defined as *n*_*i*_ and hence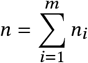, where *m* is the number of batches.

### 2.2 Phase 1: computing correlation matrix with batch effects corrected

We commence our analysis by examining the within-batch relationships. Based on the assumption that the cell sub-types are roughly evenly distributed among batches, we assume the proportion of each cell sub-type are similar. Consequently, it follows that the proportions of strong relationships (within the same sub-type) and weak relationships (between different sub-types) among all within-batch relationships should be similar across batches. From the perspective of correlation coefficients, the distributions of the within-batch correlation coefficients from −1 to 1 are expected to be similar across bathes. Note that this distribution will not be affected by batch effects as long as the assumption of even distribution holds true.

Then we consider the relationship between batches. Due to the the impact of batch effects, cells with the same sub-type but in different batches may produce undesired low correlation. Assuming that batch effect is eliminated, the correlation coefficients of samples between batches should follow a similar distribution as the within-batch correlation. Hence, by directly aligning the between-batch correlation coefficients with the distribution of within-batch correlation coefficients, we can mitigate the impact of batch effects on the correlation matrix.

In this case, the algorithm of calculating a batch-effect-free correlation matrix should be clear. We first adopt the interpolating vector quantile normalization [13] to normalize vector *v*_1_ to the distribution of vector *v*_2_, denoted as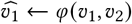. The part in the correlation matrix that corresponds to the largest batch is chosen as a reference block to obtain the reference distribution *re f*. The whole correlation matrix is hence corrected by following steps:

1. **Block-wise normalization:** Represent all non-reference within-batch blocks as a vector *w*_*i*_, then perform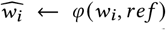. Represent all between-batch blocks as a vector *v*_*i j*_, where *i* and *j* index batches, then perform 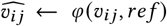 for all *i* ≠ *j*.
2. **Cell-wise normalization:** Represent all between-batch columns as a vector *v*_*i*_, where *i* index cells, then perform 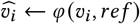 for all 1 ≤ *i* ≤ *n*.
3. **Symmetrization:** Denote the result of the last step as *D*, then the corrected matrix is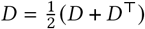.

It should be noted that while the aforementioned method is recommended for its high efficacy and efficiency, any correction method applied to the correlation matrix is deemed acceptable, as a corrected correlation matrix is the only thing we require for Phase 2 in fast-scBatch.

### 2.3 Phase 2: obtaining corrected count matrix

#### 2.3.1 Main algorithm

Through Phase 1, the corrected correlation matrix *D* is calculated. Then we expect to derive a corrected count matrix *Y* aligned with *D*.

We adopt Principal Component Analysis (PCA) to reduce the dimension in each batch. With *k* principal components each in *m* batches, a total of *mk* components are selected and combined as a reduced matrix *X*_0_ with dimension *p* × *mk*. Our goal is to obtain *Y* = *X*_0_*W* where *W* is an unknown *mk* × *n* weight matrix. Subsequently, we derived the objective function

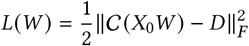

where ∥·∥_*F*_ is the Frobenius norm and C is the Pearson correlation function.

We use a gradient descent method to minimize the objective function *L*(*W*) until convergence, through a single-layer neural network without activation function. We used the Adam optimzer with adaptive learning rate. The initial value of *W* is chosen by the principal component scores within each batch, setting cross-batch initial weights to zero.

#### 2.3.2 Supplementary Algorithms

To address the challenge posed by a large number of variables (genes) in high-throughput profiling experiments, we have developed supplementary algorithms aimed at reducing computational time which may not necessarily enhancing accuracy. These algorithms are intended to be employed prior to the main algorithm, serving as rough pre-processing steps that expedite the overall process.

##### Clustering and representative cells

Cells of the same type within a batch are likely to cluster together, and their batch effects tend to exhibit similar distributions. By applying a clustering algorithm before the main procedure, we can group similar cells within each batch, resulting in clusters that contain only a few cell types. Since the batch effects within these clusters are almost identical, we can select one or a few representative cells to approximate the batch effects for the entire cluster. This strategy not only reduces the time complexity for calculating the gradient, but also accelerates the convergence.

##### Sub-sampling of genes

Given that relatively few variables dominates the principal components, considering only a random subset of components in each step of gradient descent becomes practical for accelerating the training process. Specifically, during each iteration, a subset of genes (*p*^′^ *< p*) is randomly selected, and the algorithm calculates the Pearson correlation matrix using only these genes. This method also reduces the risk of overfitting by not relying on the entire dataset in each step.

##### Deep Neural Network

Existing studies showed that the feasibility of introducing deep networks to such two-stage approach is uncertain [12]. Therefore, we also explore the effect o f deep network structures in our approach. The original algorithm use a single weight matrix *W* to adjust the raw matrix. Viewing the such approach as a single linear fully-connected layer, we attemp to extend it into a deep learning model by adding more fully connected layers and activation functions. However, in practice, this deep network approach did not perform well, suggesting that the increased complexity did not translate into better correction of batch effects.

By setting the hyper-parameters *T*_1_, *T*_2_ and *T*_3_ which controls the number of epochs in each stage, balance between efficiency and accuracy based on the specific dataset can be controlled.In summary, we propose the fast-scBatch algorithm (Algorithm 1). The workflow of the algorithm is shown in Figure 1.

**Figure 1:**
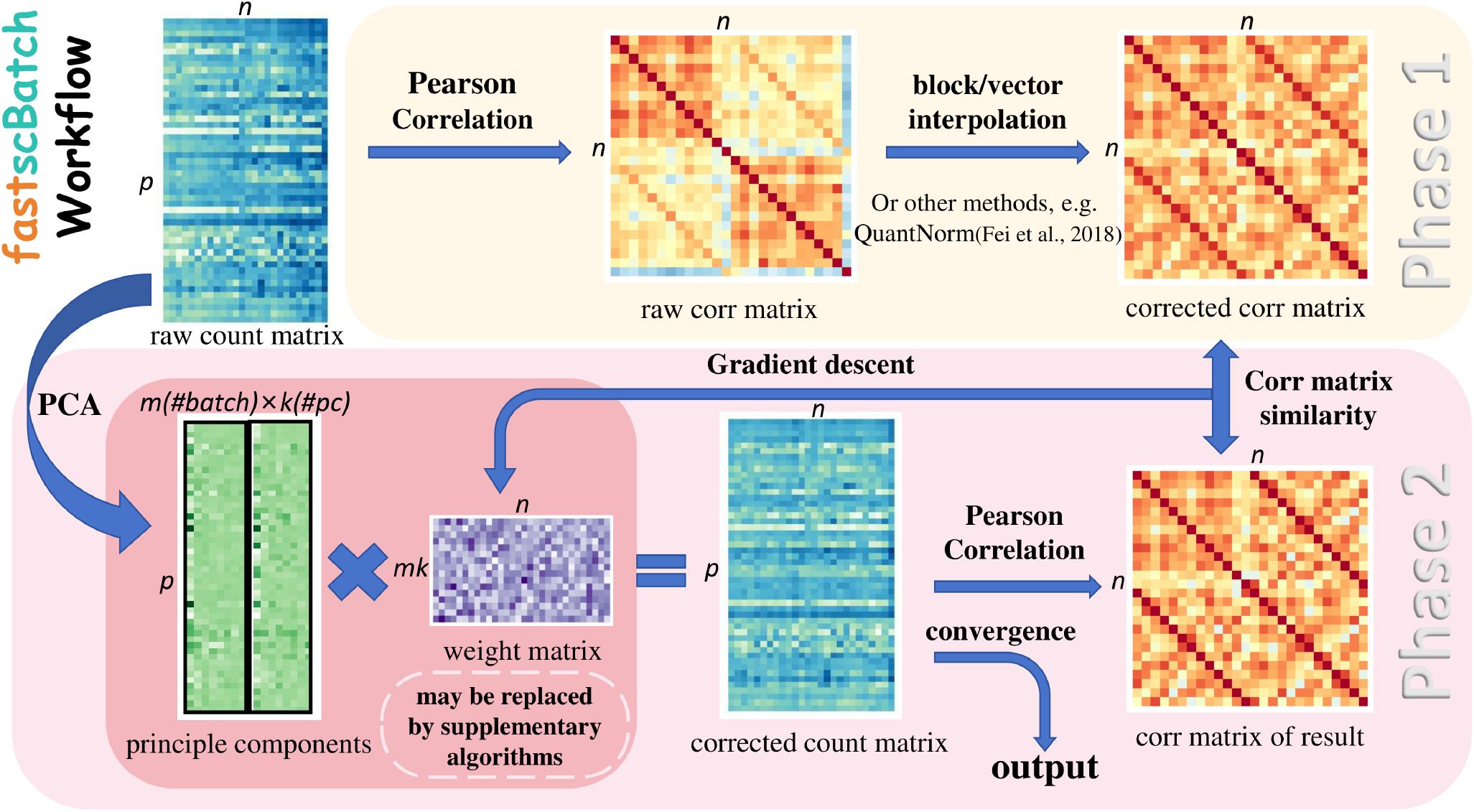
Workflow of the fast-scBatch Method. Phase 1: The raw count matrix is utilized to compute a raw correlation matrix, and then block/vector interpolation and QuantNorm are used to compute a correlation matrix with batch effects eliminated. Phase 2: The corrected matrix is given by the product of the principle components and a weight matrix, then gradient descent on the weight matrix is performed until convergence based on the similarity of the current and targeted correlation matrices.

###### Algorithm 1 Fast-scBatch Algorithm

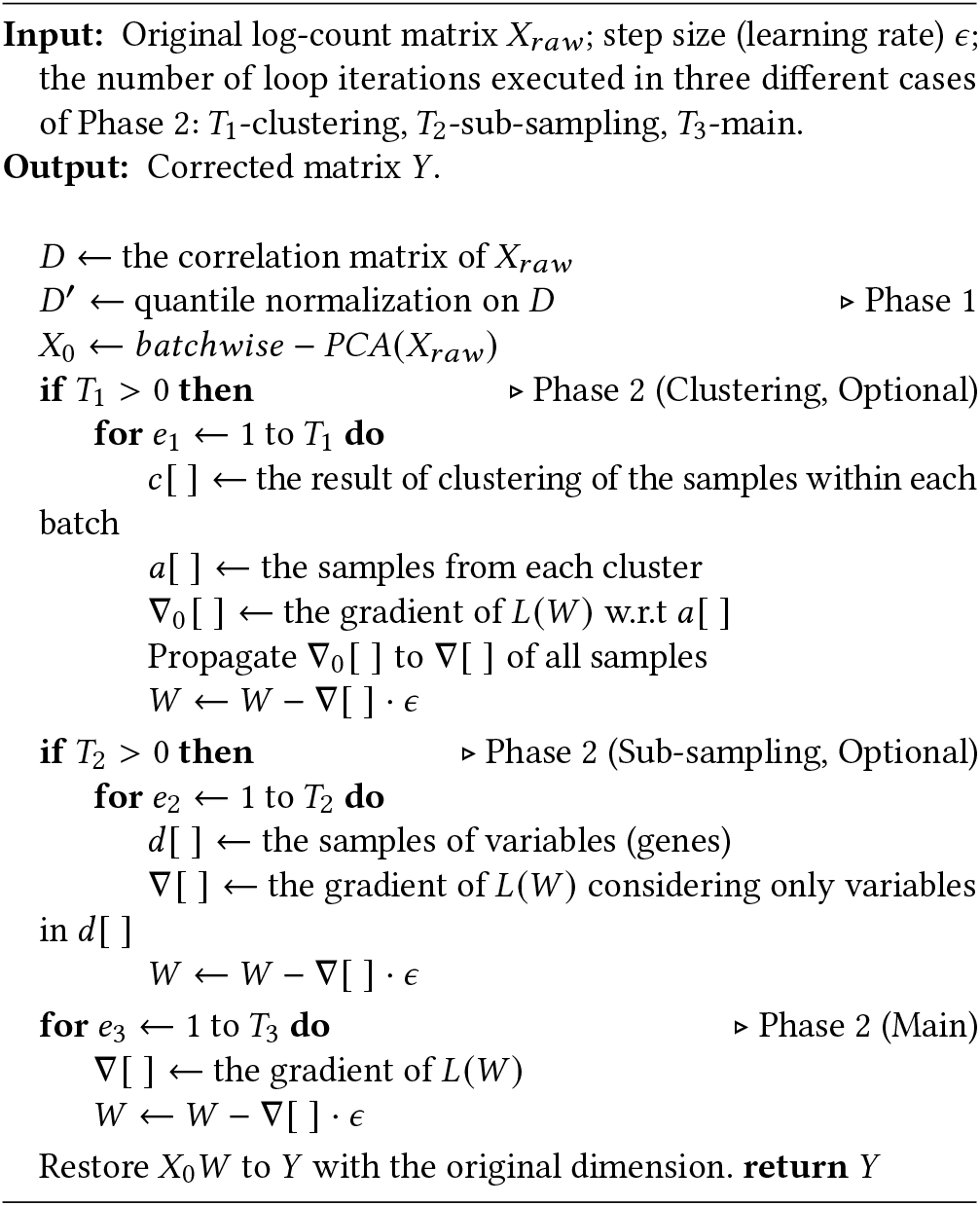

## 3 RESULTS

### 3.1 Simulation study

We utilized the R package splatter [15] to generate simulated data. With precise control over batch effects and cell type distributions, splatter facilitates a robust evaluation of various batch correction methods.

To evaluate the performance of different batch effect correction methods, we employed three sets of simulated data with sample sizes of 360, 600, and 960 cells. In each experiment, the cells were divided into four batches with proportions of 4:3:3:2. The batch effects were automatically generated by the splatter package with random parameters. These datasets enabled us to assess the scalability and effectiveness of each method as the sample size increased. The cell types in the datasets were distinguished by approximately 5% deferentially expressed genes (DE-genes), and the proportions for each cell type were 4:3:2:1.

We used several other methods for comparison, including scBatch [12], ComBat [3], MNN [6], Limma [4], and scDML[11]. Figure 2(a) illustrates the UMAP visualizations of the results in one typical simulation. In the top row, cells are color-coded by batch. In the second row, cells are color coded by cell type. We can clearly see that fast-scBatch was able to separate cell types more effectively after batch-effect correction.

To quantitatively evaluate the performance of the methods, we employed the Adjusted Rand Index (ARI) and Adjusted Mutual Information (AMI) metrics. Their definitions are as follows:

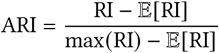

where RI is the Rand index, E[RI] is the expected value of the Rand index and max(RI) is the maximum value of the Rand index.

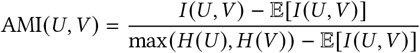

where I(U, V) is the mutual information between clustering U and V, E[*I* (*U, V*)] is the expected mutual information between U and V, *H* (*U*) and *H* (*V*) are the entropy of the clustering U and V, respectively.

These two metrics are particularly effective for measuring the accuracy of cell type identification post batch correction, as they can reflect the clustering accuracy of similar cell types from two perspectives (Figure 2(b)). The boxplot clearly illustrates that fast-scBatch deliver superior performance in both ARI and AMI metrics, showcasing its robustness in batch correction across varying sample sizes. ComBat and MNN also demonstrate commendable performance but with less consistency compared to the scBatch methods. Conversely, Limma and scDML, along with the Raw Log Count control, consistently exhibit subpar performance, highlighting their ineffectiveness in batch effect correction in the simulated setting. Considering that differences between cell types are minimized, the boxplot results reveal that some methods struggle under conditions of small sample sizes and limited information. Notably, about half of the results from linear correction methods, such as ComBat and Limma, show variability in clustering performance, a challenge that diminishes as sample size increases. On the other hand, methods that correct the correlation matrix, like scBatch and fast-scBatch, do not face this issue and maintain consistent performance across different sample sizes. Moreover, fast-scBatch outperforms scBatch in each scenario.

**Figure 2:**
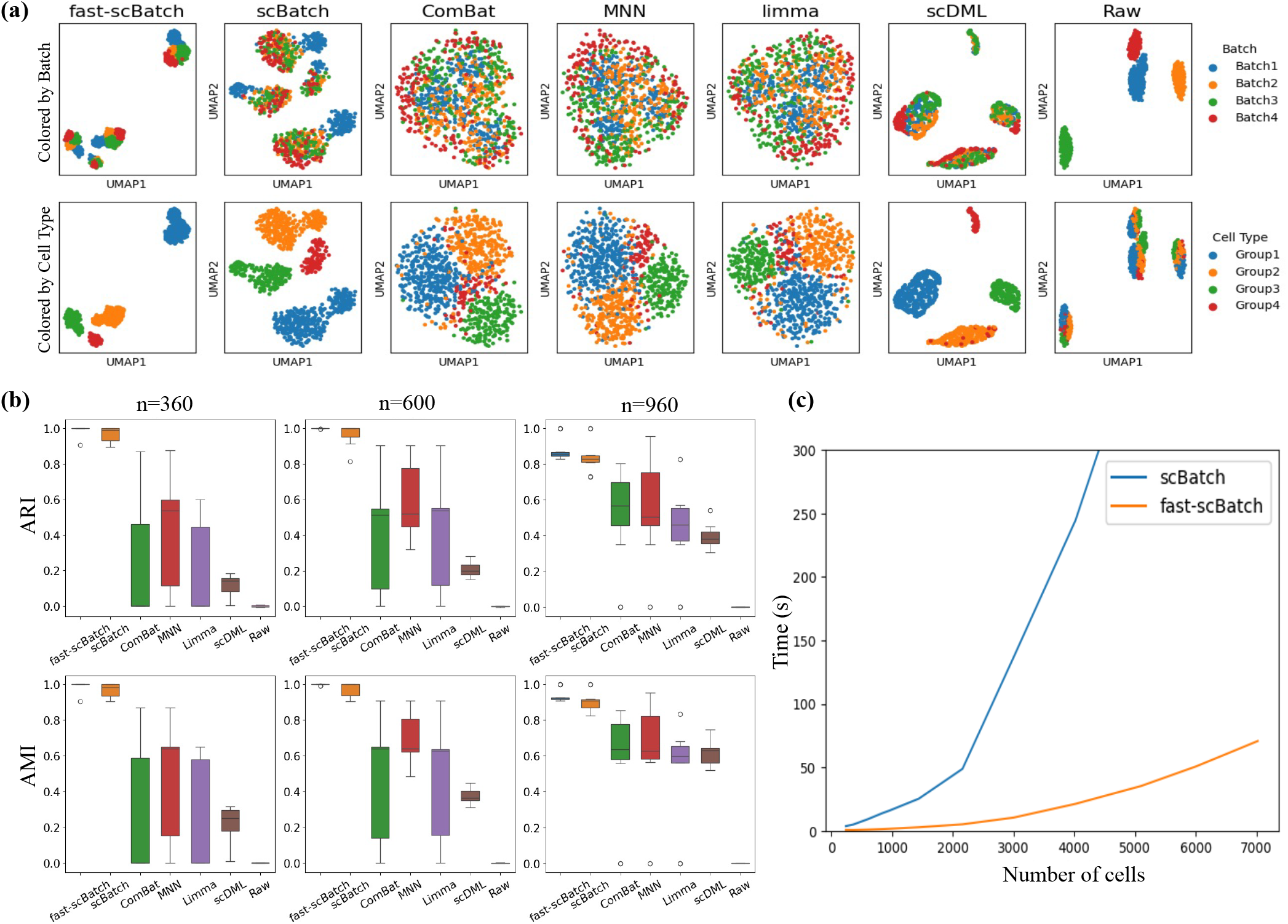
Simulation Results. (a) UMAP visualization of the simulation results, with color-coding to represent different batches and cell types. (b) Boxplot comparison of the Adjusted Rand Index (ARI) and Adjusted Mutual Information (AMI) across various methods. (c) Comparative analysis of time efficiency between scBatch and fast-scBatch, with the sample sizes ranging from 100 to 7000.

Since scBatch and fast-scBatch utilize the same approach in Phase 1 of the two-phase framework, we compared the two methods on the efficiency of Phase 2 computation. To ensure a fair comparison, scBatch was re-implemented in PyTorch, and both methods were executed on the same hardware platform. This approach eliminated any discrepancies due to differences in programming languages (R versus Python) and allowed us to focus exclusively on the inherent time complexity of each method. The computational time for the second phase of scBatch increases substantially when the sample size exceeds 2,000, becoming impractically slow for sample sizes of 5,000 or more (Figure 2(c)). In contrast, fast-scBatch demonstrates a smoother and considerably lower increase in time cost compared to scBatch. It is important to note that our re-implementation of scBatch, though based on the same computational framework and involving a comparable amount of computations, was conducted solely to ensure a fair comparison. As a result, the findings might not reflect the complete accuracy. The original scBatch algorithm, implemented in R, is even slower, struggling to process datasets with hundreds of samples within a reasonable timeframe. Therefore, from the perspective of time efficiency, fast-scBatch offers a significant improvement.

### 3.2 scRNA-seq datasets

We selected real datasets with diverse characteristics to thoroughly evaluate our methods across various scenarios. The first dataset is characterized by pronounced batch effects that overshadow sample differences, causing significant separation between cell types. Here, the primary metric for evaluation is the completeness of batch integration, which assesses the overall effectiveness of these methods. While batch effects are less pronounced in the second dataset, they are still non-negligible. Additionally, many cell types in this dataset exhibit significant similarities, potentially leading to confusion in cell type classification. Consequently, our focus shifts to the clustering accuracy of each cell type post batch effect correction, allowing us to determine whether the methods effectively preserve the original cell information without overfitting.

#### 3.2.1 Mouse neuron dataset GSE59739

The mouse neuron dataset contains four different libraries with significant batch effect [16]. We evaluate the performance of all methods by calculating the accuracy in distinguishing cells of different types (Figure 3 and Figure 4). In addition, the sizes of batches are unevenly distributed in this datasets. Despite our method’s assumption of relatively evenly distributed datasets, it performs robustly and obtains high accuracy in differentiating cell types.

**Figure 3:**
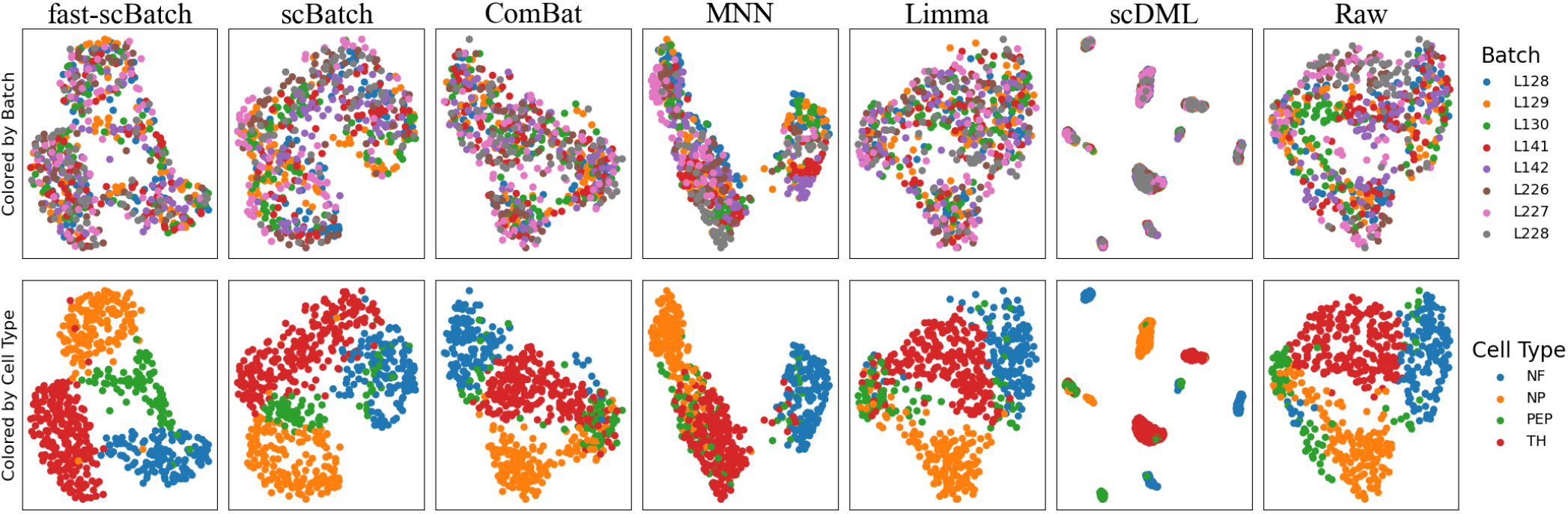
UMAP visualization of the performances for mouse neuron datasets. The first row are the results colored by batches, and the second row are the results colored by cell types. In each row, our method is plotted by the first chart from the left.

**Figure 4:**
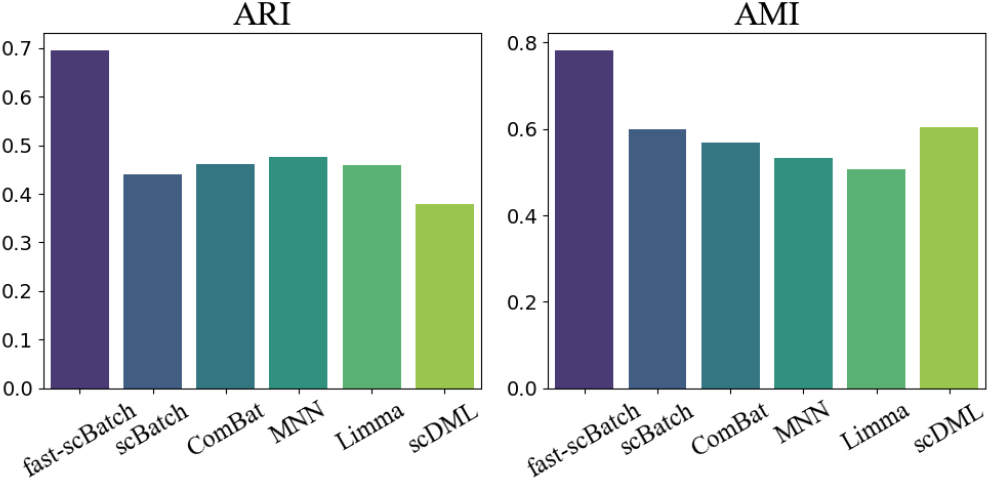
Bar-chart visualization of metrics for the mouse neuron datasets. The left panel represents the Adjusted Rand Index (ARI), while the right panel represents the Adjusted Mutual Information (AMI). In each chart, our method is represented by the first bar from the left.

Regarding the UMAP plots, the fast-scBatch method produced a clearer sample pattern, distinguishing the four subtypes more effectively. While most methods correctly merged cells of the same cell type from most or all batches, many methods can not produce a clear separation of different cell types. For example, MNN successfully integrated the PEP cells across batches, as indicated by the clustering of green samples in the UMAP plot. However, it failed to properly separate other cell types, such as NP cells (in orange), making it challenging to distinguish these types using standard clustering algorithms. Besides, methods such as scDML produces tighter clustering patterns, yet the batches are not well integrated. Fast-scBatch not only accurately integrated all batches, but also exhibits clear boundaries for all clusters, as reflected in the performance metrics (ARI = 0.69, AMI = 0.78), which are significantly higher than other methods. Notably, fast-scBatch preserved the cell type information from the uncorrected data with high contrast, a quality that other approaches failed to maintain as effectively.

#### 3.2.2 Human pancreas dataset GSE84133

The human pancreas dataset utilized inDrop single-cell RNA sequencing (scRNA-seq) to analyze over 12,000 pancreatic cells from four human donors and two strains of mice [17]. Initially, we compared the differences between human and murine cells, discovering significant discrepancies in both cell types and their quantities within the datasets. Consequently, we decided to exclude the mouse cell data from further analysis.

Our subsequent focus was on the human pancreatic cells. Despite the expectation that these cells from the four donors would exhibit similar distributions, batch effects were observed, undermining this assumption. Therefore, we concentrated on correcting the human pancreatic cell data. To evaluate the effectiveness of the correction methods, we employed the UMAP plot (Figure 5) as well as Adjusted Rand Index (ARI) and Adjusted Mutual Information (AMI) metrics (Figure 6). We note that due to the large amount of cell sample, scBatch can not produce results in reasonable time. Hence it is excluded in the comparison.

**Figure 5:**
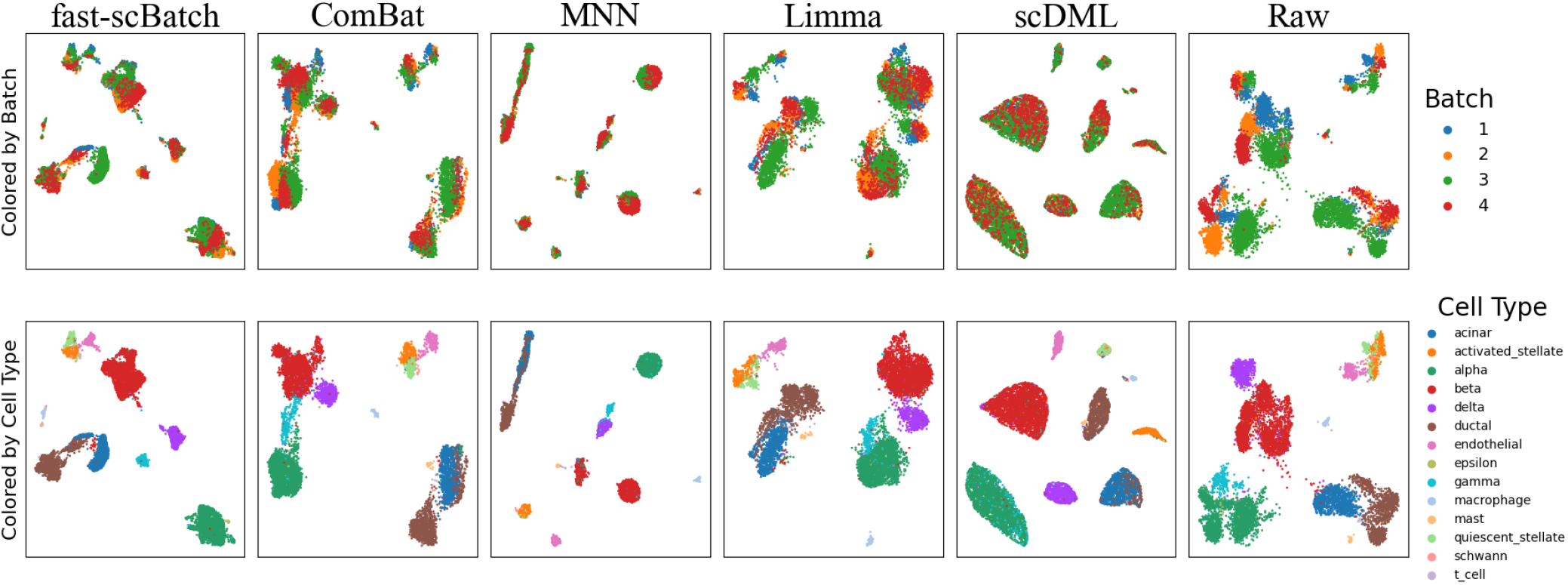
UMAP visualization of the performances for human pancreas datasets. The first row are the results colored by batches, and the second row are the results colored by cell types. In each row, our method is plotted by the first chart from the left.

**Figure 6:**
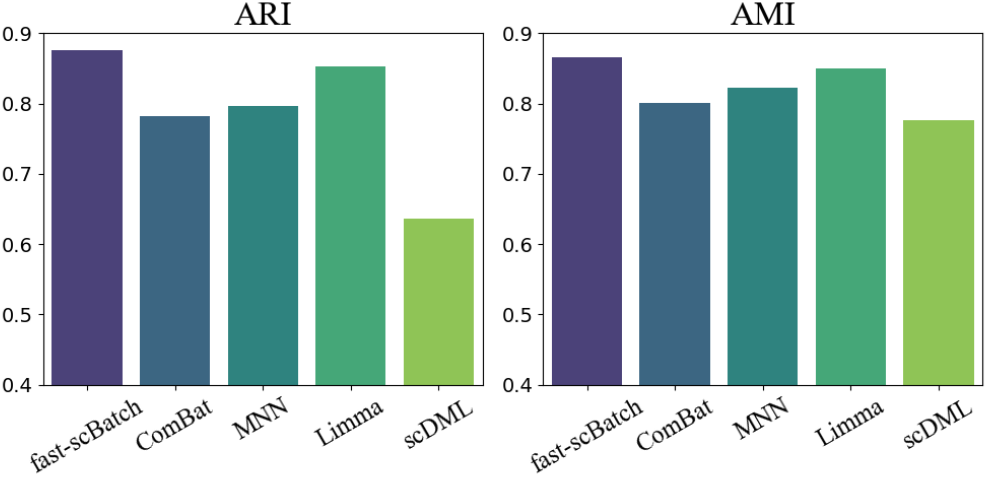
Bar-chart visualization of metrics for the human pancreas datasets. The left panel represents the Adjusted Rand Index (ARI), while the right panel represents the Adjusted Mutual Information (AMI). In each chart, our method is represented by the first bar from the left.

In the human pancreas datasets, the differences among cell types are more pronounced than the batch effect among samples. Consequently, cells of the same type still tend to cluster together. However, the significant batch effects that separate cells of the same type cannot be ignored, resulting in multiple clusters for each cell type. Hence, it is reasonable for all methods to exhibit good performance in merging the batches. However, the subtle differences among the cell types present a greater challenge. Given that many cell types share large similarities, it is difficult for any method to fully and accurately distinguish all cell types after correcting batch effects. The UMAP analysis shows that certain methods, such as MNN and scDML, are less effective at accurately clustering the cells with same cell types, resulting in lower ARI and AMI scores compared to other methods. This result indicates that these methods may lose necessary information during the correction. On the other hand, methods such as scBatch and Limma achieve higher cell-type clustering metric scores, which shows that they preserve abundant cell information though the batch are not fully integrated as shown in the UMAP plot.

Our method, fast-scBatch, demonstrates a balanced performance. It achieves slightly higher ARI scores than other methods while also producing a well-integrated UMAP plot, which means it preserves crucial cell information and also fully eliminates the batch effects. Consequently, fast-scBatch proves to be more effective in handling such sort of datasets.

## 4 DISCUSSION

Batch effect elimination is crucial for ensuring the accuracy and reliability of downstream analysis in high-throughput experiments. Uncorrected batch effects can introduce biases and confound results, leading to erroneous conclusions and impeding scientific discoveries.

Our study indicates that the proposed method, fast-scBatch, demonstrates robust and accurate performance in complex scenarios, including those with non-linear batch effects and unevenly distributed cell types. Furthermore, fast-scBatch shows promise in achieving efficient computational speed when compared to other algorithms utilizing the same two-phase approach through our analysis of the UMAP plots and metrics results.

It is curious to note that even though we used dimension reduction, which might lead to some loss of information, our correction performance measured by ARI and AMI metrics is higher than similar methods without dimension reduction such as scBatch when trained for the same number of epochs (with even lower costs in terms of time). There is a possible reason for this result. When we compare the numerical results (corrected count matrices) from the two algorithms, we find that the absolute values of the expression levels, as well as the differences in mean expression levels for each cell type, are larger in fast-scBatch compared to linear correction methods including ComBat and other two-phase methods including scBatch. This indicates that fast-scBatch, through the principle component analysis, can identify the most important genes or gene groups more accurately. Consequently, during the gradient descent process, more weight is given to these principal components. It can be concluded that PCA captures the main factors causing batch effects, allowing fast-scBatch to focus on correcting them more effectively. In addition, since we always keep a reference vector or block when correcting the correlation matrix in Phase 1, the interrelationship of the expression levels for the batch with the most samples remains relatively unchanged. This approach ensures that the structure of the largest batch serves as a stable reference, making the final results reliable as long as fast-scBatch correctly clusters the same cell types. Hence, although we correct the genes in the principal components sharply, overfitting or distortion of the datasets is unlikely to occur. Therefore, although performing PCA on the raw count matrices might lose some information, fast-scBatch can still perform better by accurately targeting and correcting the key sources of batch effects while preserving the integrity of the data. This explains why fast-scBatch shows better results despite the potential drawbacks of dimension reduction.

Another observation is that, compared to scBatch, although the primary focus of fast-scBatch was on accelerating computational speed for large datasets, it also achieved more stable performance when the sample size was small, as evidenced in the simulation result, where the range of the box-plot are smaller for fast-scBatch than other methods. This stability is likely due to the ability of principal components to effectively capture the overall effects of the batch effects, whereas the entire dataset might provide misleading information due to random factors. Smaller datasets are more susceptible to such random variations. By focusing on principal components, methods incorporating dimension reduction such as our method and scDML mitigates these issues and maintains reliable performance in small sample settings. In addition, our method incorporates some random factors in both phases to provide a general and fast solution.

There is still room to improve our method. The first concern is that, although our method demonstrates robustness to unevenly distributed datasets to some extent, it may not always produce good results when the cell type distribution in each batch becomes extremely unbalanced, as we primarily assume a relatively even distribution of cell types across the datasets at the beginning.

The second area for improvement is that our method necessitates the calculation of a corrected *n* × *n* correlation matrix in Phase 1. While low-level optimizations and enhanced hardware can marginally reduce the time cost, this approach is not memory efficient. Currently, our method supports the correction of around 20, 000 samples within minutes on GPUs, but it requires at least 8GiB of memory to store both the raw data and the correlation matrix, which represents a maximum capacity for many common single GPUs. With the increasing size of datasets, calculating and storing the entire correlation matrix may become more challenging, posing obstacles to our current algorithm. As this is a general issue affecting many methods in this field, future advancements such as GPU parallel computing are expected to benefit all methods.

Finally, although we utilize PCA to accelerate the method, which has proven effective in many cases, some complex cell distribution patterns are difficult or unsuitable to summarize with a few principal components, potentially resulting in inaccurate outcomes. It is recommended to visualize the dataset and evaluate whether the patterns are suitable for our method prior to its application.

## 5 CONCLUSION

In this study, we introduced fast-scBatch, a novel method for correcting batch effects in single-cell RNA sequencing (scRNA-seq) data. Our approach, based on a two-phase framework, integrates correlation matrix correction, within-batch principal component analysis (PCA), and gradient descent to enhance computational efficiency and maintain accuracy. Through comprehensive simulation studies using the R package splatter and evaluations on real scRNA-seq datasets, we demonstrated that fast-scBatch outperforms many existing methods in some scenarios in terms of ARI and AMI metrics for cell type identification and UMAP analyses.

Our findings highlight the robustness of fast-scBatch in handling complex cases, such as non-linear batch effects and unevendistributed cell types. Despite the potential information loss associated with dimension reduction, fast-scBatch consistently delivered higher performance.Additionally, the method’s efficient computational speed makes it particularly suitable for large datasets, addressing the growing demand for scalable batch correction solutions. Though fast-scBatch has shown its excellent performance, we should not ignore certain limitation, including the weakness in highly imbalanced datasets, the high memory occupation in calculating the correlation matrix and potential information loss through PCA.

Overall, fast-scBatch represents a significant advancement in batch effect correction for scRNA-seq data, offering a balance between speed and accuracy. Future work will focus on further optimizing the algorithm, exploring more advanced computational techniques, and extending its applicability to a wider range of datasets and experimental conditions.

## DATA AVAILABILITY

The scripts for conducting simulation experiments and real datasets examinations can be found in the Github repository https://github.com/frankchenfu/fast-scBatch-paper-script.

## ACKNOWLEDGMENTS

This research has been supported by the National Key R&D Program of China (2022ZD0116004), Guangdong Talent Program (2021CX02Y145), Guangdong Provincial Key Laboratory of Big Data Computing, and Shenzhen Key Laboratory of Cross-Modal Cognitive Computing (ZDSYS20230626091302006).

